# Sex-specific body mass ageing trajectories in adult Asian elephants

**DOI:** 10.1101/2020.12.17.423208

**Authors:** Lucas D. Lalande, Virpi Lummaa, Htoo H. Aung, Win Htut, U. Kyaw Nyein, Vérane Berger, Michael Briga

## Abstract

In species with marked sexual dimorphism and where one sex undergoes stronger intrasexual competition, that sex is expected to age earlier or quicker. Here, we utilise a unique, longitudinal dataset of a semi-captive population of Asian elephants (*Elephas maximus*), a species with marked male-biased intrasexual competition, with males being larger and living shorter, and test the hypothesis that males show earlier and/or faster body mass ageing than females. We show sex-specific body mass ageing trajectories: adult males gained weight up to the age of 48 years old, followed by a decrease in body mass until natural death. In contrast, adult females gained body mass with age until a body mass decline in the last year of life. Our study shows that sex-specific life-histories shape ageing patterns, consistent with the predictions of the classical theory of ageing.

## Introduction

Ageing – a decline in organismal functioning with age (Monaghan et al., 2008) – has been observed in many species (Jones et al., 2014). However, the onset and rates of ageing differ both between (Jones et al., 2014) and within species (Nussey et al., 2007) and between sexes (Clutton-Brock & Isvaran, 2007; Lemaître et al., 2020; Tidière et al., 2015). A main challenge in ageing research is to quantify and explain such differences in the onset and rates of ageing (Rando & Wyss-Coray, 2021).

In species with sex-specific intrasexual competition, classical theory of ageing predicts that the sex with the highest intrasexual competition has a shorter lifespan and an earlier onset and/or higher rate of ageing (Williams, 1957). The rationale is that high intrasexual selection often results in one sex showing conspicuous displays or aggressive intrasexual behaviours, leading to increased mortality and a *live fast, die young* pace of life (Bonduriansky et al., 2008; Clutton-Brock & Isvaran, 2007; Maklakov & Lummaa, 2013). For that sex, antagonistically pleiotropic genes or deleterious mutations are not counter selected due to a weakened force of selection in late-life (Williams, 1957). Accordingly, in polygynous species with male-biased intrasexual competition, males often die earlier (Lemaître et al., 2020) and age earlier or faster than females (Beirne et al., 2015; Clutton-Brock & Isvaran, 2007; Douhard et al., 2017; Nussey et al., 2009; Tidière et al., 2015). However, recent conceptual developments have shown that this association can be disrupted. This can occur for example because of condition-dependent extrinsic mortality selecting particularly high-performing individuals in the population (Chen & Maklakov, 2014) or canalisation (*i.e.* the more a trait contributes to fitness, the less it should deviate from optimal trait value, with respect to environmental variation (Flatt, 2005)), thereby contradicting the theoretically expected earlier or faster ageing in males. The extent to which such phenomena occur in nature remains unknown.

Here, we used a unique long-term dataset to describe sex-specific body mass ageing trajectories in a nutritionally unsupplemented semi-captive timber population of Asian elephants (*Elephas maximus*) living in their natural environment in Myanmar. Body mass is of interest in the study of ageing because it is positively associated with key life-history traits such as reproduction and lifespan in many non-human species (Briga et al., 2019; Gaillard et al., 2000; Hämäläinen et al., 2014; Pelletier et al., 2007). Therefore, the study of body mass ageing fits into the evolutionary framework of ageing. Accordingly, in Asian elephants, seasonal variation in body mass was positively associated with survival the following month (*e.g*. low body mass was associated with low survival during dry season) (Mumby, Mar, Thitaram, et al., 2015). Moreover, male Asian elephants benefit from being heavy during intrasexual competition for dominance and mating (Sukumar, 2003).

However, we know almost nothing about body mass ageing in elephants despite the interest in studying ageing in such a long-lived, social and sexually dimorphic non-human species. While females live in kin groups, adult males often roam solitarily, undergo a more intense intrasexual competition for dominance and mating (Sukumar, 2003) and hence are bigger, heavier (Mumby, Chapman, et al., 2015), more aggressive and less sociable (Seltmann et al., 2019) and shorter-lived than females (respective median lifespans in this population: 30.8 and 44.7 years) (Lahdenperä et al., 2018). Based on this male-biased intrasexual competition and shorter lifespan, and following the classical theory of ageing (Williams, 1957), we tested the prediction that males experience an earlier and/or faster body mass loss than females (Bonduriansky et al., 2008; Maklakov & Lummaa, 2013).

## Material and methods

### Study population

We studied the world’s largest semi-captive Asian elephant population consisting of around 3,000 individually-marked elephants owned by the government-run Myanma Timber Enterprise (MTE) (Leimgruber et al., 2008). Their birth, death, maternal-lineage pedigree, and morphological measurements have been recorded for almost a century by local veterinarians. These elephants are distributed across Myanmar in forest camps and used as riding, transport and drafting animals. Elephants work during the day and, at night, they socialise, mate and forage freely, unsupervised in forests (Oo, 2010; Zaw, 1997). There are no husbandry procedures and timber elephants are never culled. Calves born in captivity are cared for and nursed by their biological mother and allomothers (Lahdenperä et al., 2016; Lynch et al., 2019). Therefore, breeding rates are natural with no reproductive management. Moreover, there is minimal food provisioning, but elephants benefit from veterinary care that consists of treatment of simple injuries and monitoring of working conditions.

Both males and females are used in the workforce, and each working group of six elephants is composed of both sexes. Males and females follow the same government set limitations on taming age, working and retirement age, working-days per year, hours of work per day and tonnage pulled annually apply to both sexes, although it is possible that males might be used for somewhat different working tasks at times (*e.g*. when use of tusks is required; only males can possess long tusks in Asian elephants). Pregnant females are given a rest period from mid-pregnancy (around 11 months into gestation) until the calf is 1-year-old (Toke Gale, 1974), while they and their calf are being monitored by their mahouts (individual caretakers and riders) throughout this period. Following this break, mothers are used for light work but are kept with calves at heel and able to suckle on demand until the calf is four or five years old (Oo, 2010) at which point calves are assigned a rider, name, logbook and registration number. After the training period, elephants are used for light work duties until the age of 17, when they enter the full workforce until retirement around age 50. The MTE maintains their care and logbooks until death.

### Data collection and selection

Our analyses focused on age- and sex-specific variation in adult body mass from age 18 onwards, in order to omit the phase during which elephants grow in height (Mumby, Chapman, et al., 2015) and to focus only on adult body mass age-specific variations. We compiled a total of 3,886 body masses on 493 individuals (2,570 body masses on 322 females, and 1,316 body masses on 171 males). These data came from two sources: (i) body masses were either measured on elephants on the field or (ii) estimated using height to the shoulder and chest girth (method in Supplementary Information 1). For the first source, we extracted 1,901 body masses of 347 elephants (1,297 measurements on 230 females, and 604 measurements on 117 males) and for the second source we estimated 1,985 body masses on 342 individuals (1,273 estimations on 226 females, and 712 estimations on 116 males - a same individual can have both measured and estimated body masses). For all elephants of the 325 working localities (‘township’) sampled, sex, year of birth (‘YOB’), alive or dead status at the moment of the study, origin (captive-born or wild-caught) and measurement season (hot: Feb-May, monsoon: Jun-Sep, cold: Oct-Jan (Mumby, Mar, Thitaram, et al., 2015)) were known. The alive or dead status was used to test for potential terminal decline. Among the 493 individuals considered, 5 males (63 observations) and 18 females (185 observations) were known to be dead. We had measurements during the last year of life for 2 males (7 observations) and 10 females (54 observations). Study elephants were aged 18 – 72 years (mean = 39.3) and born 1941 – 1999. Age and cohort information were comparable between sexes, with 171 males (n = 1,316) born 1954 – 1999 and aged 18 – 64 years (mean = 37.4), and 322 females (n = 2,570) born 1941 – 1999 and aged 18 – 72 years (mean = 40.2).

Most elephants of this semi-captive population get at least occasionally measured for height and chest girth, with no selection with respect to their age, sex or condition. Body mass is measured only in camps provided with measurement scales (mainly in regions with the highest concentrations of elephants and the best accessibility). All elephants within the reach of those camps get weighed, again without any bias regarding their age, sex or condition. The logbooks containing these measurements have thus far been translated from Burmese to English mainly from the Sagaing region for logistic reasons, but again without any bias or pre-selection of certain individuals.

In total, we obtained a median of 4.0 measurements/individual (2.5 – 97.5^th^ percentiles: [1.0; 36.4], followed during a median period of 2.8 years (2.5 – 97.5^th^: [0.0; 36.6] on 493 elephants (n = 3,886). Two influential observations measured at age 18 and 23 were removed for one male because of particularly low Δage (Cook’s distance = 0.61 and 0.25, mean of 0.001 on all males). Other observations for this male, all after age 50, were included.

### Statistical analyses

We investigated the age- and sex-specific variation in body mass in R version 4.1.1 (R Core Team, 2021), using the body mass (log-transformed to reach normality of the variable and because of the allometric relationship between body mass and size) as a dependent variable with a normal error distribution. First, we tested in a single model, whether there were sex-specific ageing trajectories using an interaction term (Table S2). Given that this interaction was statistically significant, we compared the sex-specific ageing trajectories for both sexes separately. We did these analyses using both general additive mixed models (GAMMs) with cubic regression splines (but note that other spline functions gave similar conclusions as those shown here) and general linear mixed models (GLMMs) with respectively the functions ‘gamm’ of the package ‘mgcv’ (v. 1.8-36, Wood, 2011) and the function ‘lmer’ of the package ‘lme4’ (v. 1.1-27.1, Bates et al., 2015)). GAMMs allow more flexible ageing trajectories than GLMMs, but the more constrained ageing trajectories in GLMMs allow a less descriptive identification of differences in ageing trajectories (Fig. S1) and both approaches gave consistent conclusions (see results section). We identified the best fitting models using the model selection approach based on the second order Akaike Information Criterion (AICc) as implemented in the package ‘MuMIn’ (v. 1.43.17, Bartoń, 2021). In brief, the best fitting model has the lowest AICc value, with other models within 4 ΔAICc being plausible and models becoming increasingly equivocal up to 14 ΔAICc, after which they become implausible (Burnham et al., 2011). Visual inspection of model residuals confirmed that these fulfilled all assumptions of distribution and homogeneity without any influential data points or outliers (see above).

#### Within- vs. between-individual change

In all models, we accounted for non-independence of data due to repeated measurements from the same individual by including elephant identity (‘ID’) as a random intercept. The composition of the population can change with age for example due to selective disappearance of certain (*e.g*. lighter or heavier) individuals, which can affect the age trajectory. In order to alleviate as much as possible this problem in such a long-lived species, we decomposed body mass changes with age into between- and within-individual changes following the approach developed by van de Pol & Verhulst, (2006) and van de Pol & Wright, (2009) using two terms: *i*) the age at last measurement for each individual, which captures the between-individual variations and *ii*) a ‘Δage’ term (age at measurement minus the individual’s mean age for all measurements) capturing the within-individual changes with age. We mean-centered and standardised ‘Δage’ so that *i*) individuals measured once all get a Δage = 0 and hence contribute to the variance of the Δage intercept but not to its slope and *ii*) to avoid collinearity and to have comparable variance for Δage and Δage^2^ (Bolker, 2008; Zuur et al., 2009). Among the 493 individuals of our dataset, 105 individuals had only one measurement. We included these individuals by giving them Δage = 0 (*i.e*. mean-centered) so they do not contribute to the coefficient but do contribute to the variance along the Y axis on Δage = 0, diminishing the likelihood of a false positive, and ii) do contribute to the coefficient of the age at last measurement term, thereby avoiding a bias in the dataset from selecting only the longer-lived or most monitored individuals.

#### Testing ageing trajectories

We tested several within-individual ageing trajectories, first using GAMMs, which can provide curvilinear relationships and allow to describe trends, and using GLMMs, able to detect breaking points if necessary, by testing linear, quadratic, threshold and terminal models (Fig. S1) and we selected the ageing trajectory with the lowest AICc. For GAMMs, we identified the age at which maxima occurred based on the first-order derivative (= 0) using the function ‘fderiv’ of the package ‘gratia’ (v. 0.6.0, Simpson & Singmann, 2021). For threshold models (Fig. S1C), we followed the approaches previously developed in Briga et al., (2019) and Douhard et al., (2017). In brief, we first identified the best-fitting threshold age in a series of models, varying the threshold in the ‘Δage’ term between −35 to 22 years with intervals of one Δage (1 mean-centered Δage = 4.4 and 4.5 years for males and females respectively) and estimated the threshold and its confidence intervals using ± 4 ΔAIC age range. Then we compared the best-fitting threshold model with the other ageing trajectories. Sometimes, declines in trait value appear shortly before death (terminal decline). We coded a ‘terminal’ change (Fig. S1D) as a binomial factor for whether an individual died during the year following the measurement. We used a one-year-window to avoid a possible seasonal covariation in weight and because it was the best fitting time-window, but note that models with other time-windows gave consistent conclusions (Fig. S3).

#### Accounting for temporal and spatial variation in body mass

As body mass variation can be influenced by seasonal, spatial and within-individual factors, we tested whether body mass values were affected by (i) measured or estimated, (ii) individuals were alive or dead, (iii) captive- or wild-born and (iv) the measurement season. To this end we used a model selection approach, performing a dredge on the best-fitting ageing trajectories for each sex to test for confounding factors (Table S3). In our models, we included as random intercepts individual identity to account for the repeated measurement of the same individual. We also included ‘township’ to account for the spatial clustering of individuals across Myanmar, although actually adding township worsened the model fit in most cases (male GAMM: ΔAICc = +1.5, GLMM: ΔAICc = +1.8; female GAMM: ΔAICc = +0.6, GLMM: ΔAICc = −21.2).

## Results

At the measurements’ starting age of 18 years, males were on average 235 kg heavier than females, weighing respectively 2,541 kg [95%CI: 2,406; 2,683] and 2,306 kg [95%CI: 2,258; 2,355] and this difference was statistically significant (ΔAICc = −122.6 in a GLMM with vs without sex as a fixed effect).

We identified the elephant’s body mass ageing trajectories using general additive mixed models (GAMMs) and general linear mixed models (GLMMs) and both approaches gave consistent results. Both analyses showed that sexes have different body mass ageing trajectories (interaction term, GAMM: ΔAICc = −65.7, Fig. S4, Table S2; GLMM: ΔAICc = −47.0, Fig. 1, Table S2) and hence, we identified the ageing trajectories for both sexes separately.

**Figure 1.**
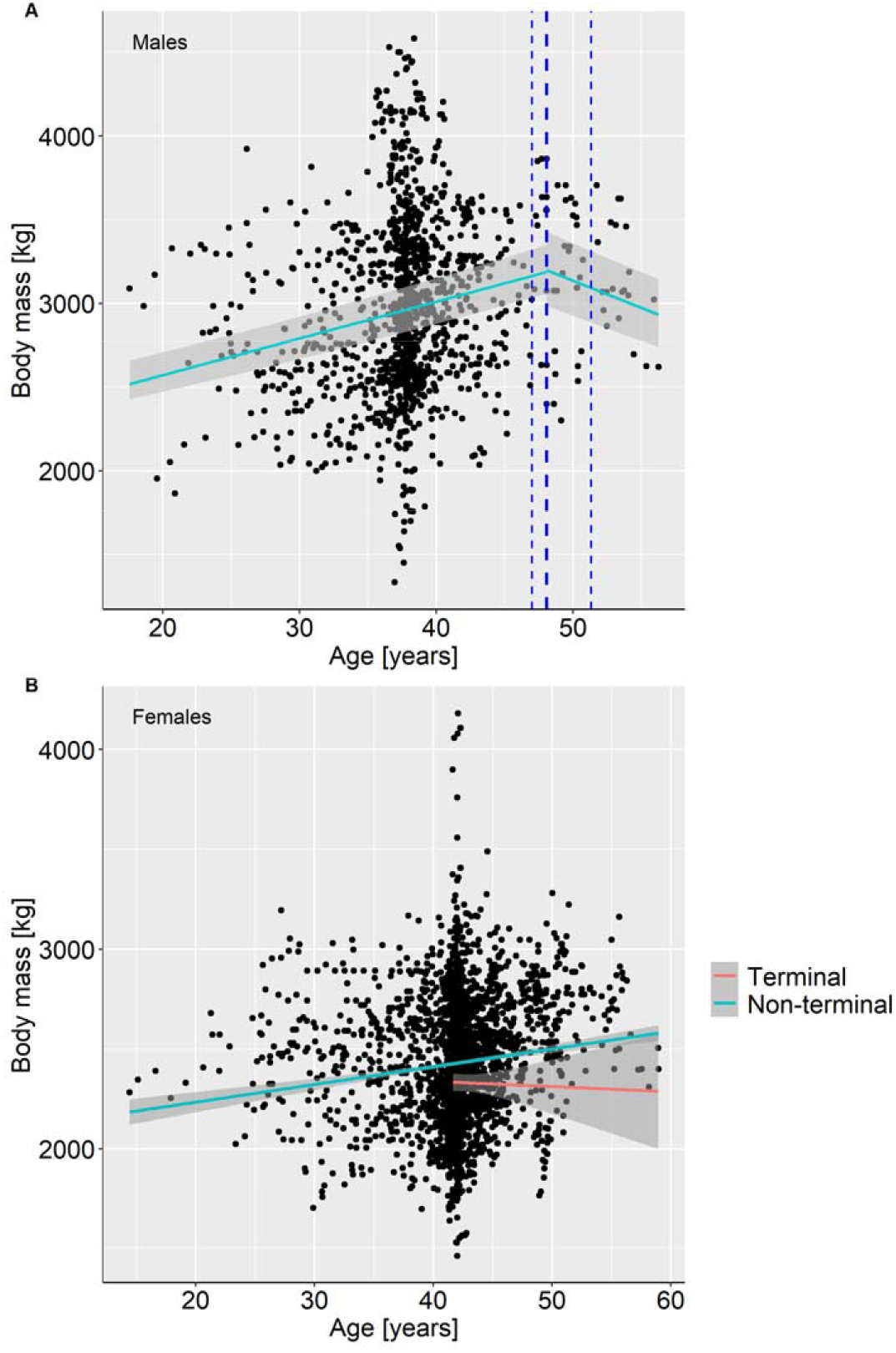
Body mass ageing trajectories of (A) males (n = 1,316 measurements on 171 individuals) and (B) females (n = 2,570 measurements on 322 individuals) with predictions of the best-fitting GLMMs (Table 1) with grey areas 95%CI. For males, the thick dashed-line shows the threshold age at onset of the body mass decline (1.9 or 48.3 years) with thin dashed-lines the 4 ΔAICc-CI [46.6, 52.3]. For females, measurements in the terminal year (red) are significantly lower (intercept) than measurements at other ages (blue). Note 1: the terminal slope is for illustration purposes only and was not statistically tested. Note 2: the original x-axis is Δage, but for simplicity, we presented here x-axis as age. For the original figure, please see Fig. S2.

For males, both GAMMs and GLMMs indicated a body mass gain from age 18 years until their late-forties or early fifties (GAMM maximum: 54 years [95%CI: 53; 56], Fig. S5; GLMM maximum: 48.2 years [4 AICc CI: 47.1; 51.6], Fig. 1A), followed by a decline until death (Fig. S4A, Fig. 1A, Table 1, Table S4). In GLMMs, this maximum was shown through the best fit of a threshold model (ΔAICc = −30.4 compared to a linear trajectory): males gained mass at a rate of 22 kg/year [95%CI: 19.4 23.7] or 1% [95%CI: 0.9; 1.1] of males’ mean body mass and then lost mass at a rate of 29 kg/year [95%CI: 14.9; 41.9] or 1.3% [95%CI: 0.7; 1.9] of males’ mean body mass (Fig. 1A, Table 2). Regarding the decline, neither GAMMs nor GLMMs showed confident statistical support that it was terminal (*i.e*. determined by time before death rather than age): in GAMMs, models with and without the terminal term were almost equivalent (ΔAICc = −0.01, Table S4) and in GLMMs adding a terminal term worsened the model fit (ΔAICc = +5.0, Table 1, Fig. S3A).

**Table 1.** Best fitting body mass ageing trajectories (bold) for males and females, using GLMMs for each model ageing trajectories ranked from the least to the most complex. AICc: second-order Akaike Information Criterion; ΔAICc: change in AICc relative to the best fitting model; k: degrees of freedom.

| Model type | Model | Males |  |  | Females |  |  |
| --- | --- | --- | --- | --- | --- | --- | --- |
| | | K | AICc | ΔAICc | k | AICc | ΔAICc |
| null | $\log(\text{bm}) \sim 1$ | 4.0 | -2,828.3 | 364.2 | 4.0 | -5,442.9 | 160.7 |
| linear | $\log(\text{bm}) \sim \Delta\text{age} + \text{age-last}$ | 6.0 | -3,162.0 | 30.4 | 6.0 | -5,586.1 | 17.5 |
| +terminal | $\log(\text{bm}) \sim \Delta\text{age} + \text{age-last} + \text{terminal}$ | 7.0 | -3,156.9 | 35.6 | 7.0 | -5,592.3 | 11.3 |
| age-last<sup>2</sup> | $\log(\text{bm}) \sim \Delta\text{age} + \text{age-last} + \text{age-last}^2$ | 7.0 | -3,155.5 | 37.0 | 7.0 | -5,598.7 | 4.9 |
| **+terminal** | **$\log(\text{bm}) \sim \Delta\text{age} + \text{age-last} + \text{age-last}^2 + \text{terminal}$** | 8.0 | -3,150.4 | 42.1 | **8.0** | **-5,603.6** | **0.0** |
| $\Delta\text{age}^2$ | $\log(\text{bm}) \sim \Delta\text{age} + \Delta\text{age}^2 + \text{age-last}$ | 7.0 | -3,176.8 | 15.7 | 7.0 | -5,571.6 | 32.0 |
| +terminal | $\log(\text{bm}) \sim \Delta\text{age} + \Delta\text{age}^2 + \text{age-last} + \text{terminal}$ | 8.0 | -3,171.2 | 21.3 | 8.0 | -5,577.9 | 25.7 |
| quadratic | $\log(\text{bm}) \sim \Delta\text{age} + \Delta\text{age}^2 + \text{age-last} + \text{age-last}^2$ | 8.0 | -3,170.1 | 22.3 | 8.0 | -5,584.3 | 19.3 |
| +terminal | $\log(\text{bm}) \sim \Delta\text{age} + \Delta\text{age}^2 + \text{age-last} + \text{age-last}^2 + \text{terminal}$ | 9.0 | -3,164.6 | 27.9 | 9.0 | -5,589.3 | 14.3 |
| **threshold** | **$\log(\text{bm}) \sim \Delta\text{age1} + \Delta\text{age2} + \text{age-last}$** | **8.0** | **-3,192.5** | **0.0** | 8.0 | -5,580.5 | 23.1 |
| +terminal | $\log(\text{bm}) \sim \Delta\text{age1} + \Delta\text{age2} + \text{age-last} + \text{terminal}$ | 9.0 | -3,187.5 | 5.0 | 9.0 | -5,586.0 | 17.6 |
| threshold (age-last<sup>2</sup>) | $\log(\text{bm}) \sim \Delta\text{age1} + \Delta\text{age2} + \text{age-last} + \text{age-last}^2$ | 9.0 | -3,185.8 | 6.6 | 9.0 | -5,592.6 | 11.0 |
| terminal | $\log(\text{bm}) \sim \Delta\text{age1} + \Delta\text{age2} + \text{age-last} + \text{age-last}^2 + \text{terminal}$ | 10.0 | -3,180.9 | 11.6 | 10.0 | -5,596.9 | 6.7 |

**Table 2.** Parameter estimates of linear mixed-effect models including individual body mass beyond 18 years of age as the response variable (in kg, log-transformed) for male and female Asian elephants. V: variance, SD: standard-deviation, SE: standard-error. Marginal and conditional R^2^ give the variance explained by fixed effects, and both fixed and random effects, respectively.

| Males |  |  | Females |  |  |
| --- | --- | --- | --- | --- | --- |
| Random effects | V | SD | Random effects | V | SD |
| Individual identity | 0.019 | 0.137 | Individual identity | 0.011 | 0.103 |
| Township | 0.0005 | 0.023 | Township | 0.003 | 0.056 |
| Fixed effects | Estimate | SE | Fixed effects | Estimate | SE |
| Intercept | 7.707 | 0.021 | Intercept | 7.840 | 0.019 |
| Age at last measurement | 0.087 | 0.012 | Age at last measurement | 0.025 | 0.007 |
| Δage1 | 0.041 | 0.002 | Age at last measurement<sup>2</sup> | -0.028 | 0.006 |
| Δage2 | -0.057 | 0.014 | Δage | 0.016 | 0.001 |
|  |  |  | Terminal (1) | -0.071 | 0.020 |
| **Marginal R<sup>2</sup>** | 0.24 |  | **Marginal R<sup>2</sup>** | 0.10 |  |
| **Conditional R<sup>2</sup>** | 0.89 |  | **Conditional R<sup>2</sup>** | 0.77 |  |

For females, both GAMMs and GLMMs indicated a body mass gain throughout their lives until a terminal decline during their last year of life (Fig. S4B, Fig. 1B, Table 1, Table S4). GLMMs indicated a mass gain of 9 kg/year [95%CI: 7.5; 10.4] or 0.35% [95%CI: 0.3; 0.4] of females’ mean body mass (ΔAICc = −6.7, Table 1, Table 2). Loss of body mass occurred in the last year of life (GAMM: ΔAICc = −12.2, Table S4, Fig. S4B; GLMM: ΔAICc = −4.9, Table 1, Fig. 1B, Fig. S3B) and consisted of 173 kg [95%CI: 80; 263] or 6.8% [95%CI: 3.2; 10.4] of their mean body mass (Table 2). For comparison, the extent of the terminal decline in males, if any, is just over half that in females at 96 kg [95%CI: −19; 205] or 4.3% [95%CI: −0.9; 9.2] of males’ mean body mass (quantified in the aforementioned best-fitting threshold model). Note that for females and for both GAMMs and GLMMs, the second best model confirms the linear age trajectory, but excludes the terminal effect (GAMM: ΔAICc = +12.2, Table S4; GLMM: ΔAICc = +4.9, Table 1). For males, the second best model (for GAMMs and GLMMs) conserves the same threshold trajectory, but includes the terminal effect (GAMM: ΔAICc = −0.01, Table S4; GLMM: ΔAICc = +5.0, Table 1, Fig. S3B). Model averaging on ageing trajectories within 7 ΔAICc (Burnham et al., 2011) confirmed the ageing trajectories found, *i.e*. a threshold trajectories for males with a significant decline of body mass from 48 years old onwards (GLMM: β = −0.06 [95%CI: −0.09; - 0.03]) and a non-significant terminal decline (GLMM: β = −0.04 [95%CI: −0.10; 0.01], GAMM: β = −0.04 [95%CI: −0.09; 0.01]). Similarly, model averaging performed on models within 7 ΔAICc for females confirmed the ageing trajectory found, *i.e.* a body mass gain throughout life (GLMM: β = 0.015 [95%CI: 0.01; 0.02]) with no significant decline before the last year of life (GLMM: β = −0.07 [95%CI: −0.11; −0.03]). Also, we found no effect of none of the temporal and spatial confounding variables tested (Table S3).

## Discussion

We tested whether in a species with marked male-biased intrasexual competition, males showed an earlier and/or faster rate of body mass ageing than females. Both sexes gained mass during early adulthood. However, the onset of body mass ageing differed between both sexes: males began to lose mass from 48.3 years old onwards. In contrast, females lost body mass generally at an older age, namely in their last year of life. Compared to a previous study on growth curves of this population (Mumby, Chapman, et al., 2015), we are now using a larger dataset, including older and more numerous retired individuals. This allowed us to evidence body mass ageing in this species, which was not possible until now. Here we discuss the implications of our results in the light of the classical theory of ageing (Williams, 1957) and of the management of Asian elephants.

We describe for the first time a sex-specific pattern of body mass ageing in this species. Body mass ageing is often used in mammals as it may underpin actuarial and reproductive ageing (Beirne et al., 2015; Bérubé et al., 1999; Nussey et al., 2011). In our population, sex-specific mortality ageing has been already shown (Lahdenperä et al., 2018) and males display higher mortality than females at all ages. However, reproductive ageing has only been described in females of this population due in part to the difficulty of recording paternity in male elephants (Hayward et al., 2014; Mumby, Mar, Hayward, et al., 2015; Robinson et al., 2012). Our results provide valuable insights on how body condition declines with age and offer another aspect of the multifaceted ageing, often referred to as a mosaic (Walker & Herndon, 2010), in this long-lived and highly social species.

Asian elephants show male-biased intra-sexual competition, with males being more aggressive (Seltmann et al., 2019), fighting more for dominance and showing higher rates of mortality at all ages than females, including during early development, as calves and during adulthood (Lahdenperä et al., 2018). In such species, classical theory of ageing predicts that males should show an earlier onset or accelerated ageing (Williams, 1957). Indeed, in several polygynous mammals, males display earlier onset or higher rates of ageing than females, suggested to be due to their stronger intrasexual competition (Clutton-Brock & Isvaran, 2007 but see also Camus et al., 2012; Tower, 2006). For example, in European badgers (*Meles meles*, Beirne et al., 2015) and Soay sheep (*Ovis aries*, Hayward et al., 2015), males systematically showed stronger or earlier body mass ageing compared to females. Conversely, in monogamous species, males and females’ onsets and rates of ageing tend to be similar (Bronikowski et al., 2011; Clutton-Brock & Isvaran, 2007; Thorley et al., 2020). Our results are consistent with those studies and with the prediction of the classical theory of ageing. Moreover, our results are inconsistent with later alternatives that suggest that the prediction of the classical theory of ageing can be disrupted by high early-life condition-dependent mortality in males (Chen & Maklakov, 2014) or by canalisation (Flatt, 2005).

However, as mentioned above, previous work on this population showed both reproductive and survival age-related decline in females (Hayward et al., 2014; Robinson et al., 2012). Hence, the ageing trajectories do not synchronise between traits in females. Empirically, this heterogeneity of ageing patterns is more the rule than the exception as found in other species (Briga & Verhulst, 2021; Hayward et al., 2015; Walker & Herndon, 2010). In our population, this mismatch can be explained by the fact that body mass is a poor predictor of reproductive success, number of offspring produced or raised up to independence (5 years old), and that no relationship between height and survival has been found in females (Crawley et al., 2017). Our results that females do not show age-dependent body mass decline combined with previous results are at odds with study on other vertebrates. Asian elephants reproduce all year long (Brown, 2014), despite living in a seasonal environment, meaning that females finance reproduction through energy stored before reproduction. This is contrary to, say roe deer (*Capreolus capreolus*), an income breeder financing reproduction concurrently to gestation as this ungulate does not store reserves (Andersen et al., 2000). In the latter case, reproductive success therefore depends on body condition and available resources, while in the former case, elephants reproduce when they have stored sufficient resources to finance gestation. This might in part explain the absence of relation between female body mass ageing and reproductive senescence, contrary to males, benefiting more from being heavy than females during intrasexual competition (Sukumar, 2003).

Our study is subject to a number of limitations when it comes down to identifying why the sexes may differ in their ageing trajectories. First, it is possible that male elephants in our timber elephant population are used more for tasks requiring strength or tusks, thereby causing an earlier onset of body mass declines in males than in females. However, both sexes fall under the same government-set workload, care and retirement regulation, except for females’ maternity leave. One substantial difference between sexes is that parental care is concentrated on females, with for example only females being given ‘parental leave’ following reproduction (Toke Gale, 1974). However, since maternity is energetically expensive and no more favourable than timber working, this is unlikely to lead to the delayed onset of body mass declines in females. An ideal test would be to analyse the effect of timber work and maternity leave on body mass dynamics.

Second, elephants have a specific dentition that consists of molar teeth that eventually wear down at the end of their lives, and pathologic malocclusions or lack of molars can lead to weight loss and death by starvation. In sexually dimorphic species of ungulates, males generally display smaller molar teeth size compared to females, relatively to body size. This results in teeth wearing down faster and depleting earlier for males than females (Carranza & Pérez□Barbería, 2007) with potential consequences on male senescence compared to females. In Asian elephants, although both sexes have the same molar dental anatomy, it is possible that the earlier onset of body mass declines in males reflects sex-specific differences in tooth wear. Indeed, in captive species, dental problems have been described well before the last year of life (Gaillard et al., 2015) and, if there is sex-specific tooth wear, this could be associated with the earlier onset of body mass ageing in males.

Third, male elephants have recurring periods of physiological “musth” throughout their adult lives, which can temporarily but profoundly impact the body mass of individual males (Eisenberg et al., 1971) thereby affecting the body mass ageing trajectory. Unfortunately, recording morphological measurements is difficult during the musth period during which males display highly aggressive behaviours, although accounting for musth would improve future analyses.

Fourth, in our study, we did not find any evidence for body mass-based selective disappearance, but, as it is often the case in long-lived species, the average longitudinal individual monitoring is short relative to the lifespan of this species (*e.g.* Global BMI Mortality Collaboration et al., 2016; Prospective Studies Collaboration, 2009), and hence we only have limited power to detect such association. It is possible that there are sex-specific dynamics of selective disappearance, but whether that is the case in Asian elephants remains to be shown. An analysis with more longitudinal data would be useful to tackle this question.

Fifth, for both sexes, it is possible that maximum body mass is set by physiological and ecological constraints as indicated by the weight growth curves found earlier in this population (Mumby, Chapman, et al., 2015). These constraints could be to some extent sex-specific, although at this point, we can only speculate as to why these constraints may drive sex-specific ageing trajectory. Finally, we found a maximum body mass in males but not in females. This sex-specific differences could be driven by the fact that male elephants benefit more than females from being heavy during intrasexual competition (Sukumar, 2003).

Another factor to take into account is that retirement occurs at around 50 years in both sexes, which likely diminishes physical exercise and allows more time for foraging, thereby continuing the weight gain. The reduced intrasexual competition in females relative to males, together with this retirement, could lead to the continued mass gain of females. One way of disentangling the effect of senescence and retirement on body mass trajectories would be to know whether muscle is lost over fat. Unfortunately, we do not have the data to know this. However, it seems that muscle function does not decrease with age in this semi-captive population in neither sex (Reichert et al., 2022). On the contrary, fat storage as measured by levels of circulating triglycerides remained constant up to adulthood, decreasing afterwards in senior elephants (from retirement onwards) in both sexes (Reichert et al., 2022). Also, all elephants officially retire at age 55, but most elephants enter pre-retirement around the age of 50 because of decreased strength. These results, taken together with the onset of body mass decline we found in males (*i.e*. 48 years old), suggest that retired individuals lose fat in both sex rather than muscle and that body mass ageing is rather a cause than a consequence of retirement in males. Nevertheless, given that elephants in the wild do not experience timber labour and retirement, we cannot exclude that the sex-specific body mass ageing trajectories could be different in a wild (non-working) population of Asian elephants compared to those found in our study.

We found that females experienced a terminal body mass decline in the last year of life. Our data contain both males and females among the oldest ages (>50), hence sex-specific terminal decline is unlikely to emerge from differences in lifespan. In European badgers, a species in which females outlive males, both sexes displayed terminal body mass declines (Beirne et al., 2015). It is possible that the sex-specific terminal declines in our study resulted from differences in power, with 5 dead males and 18 dead females. Indeed, for both males and females, the coefficient and effect size of the terminal terms were negative, but the effect size in males remained about half of that in females (Cohen’s d_males_ −0.045 [95%CI: −0.10; 0.01] = a decline of 96 kg [95%CI: −19; 205], Cohen’s d_females_ −0.071 [95%CI: −0.11; −0.03] = a decline of 173 kg [95%CI: 80; 263]). Hence, it is possible that the sex-specific terminal effect is driven by power issues and we look forward to testing that with several more years of monitoring.

Terminal declines emphasise that the chronological age is rarely a perfect estimation of the biological age which can better describe the ‘true biological state’ of an organism (Klemera & Doubal, 2006). In that sense, terminal decline is a biomarker of health and remaining lifespan. The ‘terminal illness’ hypothesis refers to the age-independent decrease of a trait value, related to the imminent death of the individual (Coulson & Fairweather, 2001). Such terminal effects were shown for example for body mass in mammals (stronger in males than females in European badgers (Beirne et al., 2015), in both sexes in Soay sheep (Hayward et al., 2015) and in male but not female Alpine marmots (Tafani et al., 2013)) and for sexual signals in birds (Simons et al., 2016). For which traits or under which conditions to expect terminal declines remains yet poorly understood but our study highlights the importance of studying sex-specific differences in ageing and illustrates the need to improve our understanding of the mechanisms driving the diversity of ageing patterns in the wild.

## Acknowledgements

We thank the Myanma Timber Enterprise (MTE) and the Myanmar Ministry of Natural Resources and Environmental Conservation for their collaboration and all of the MTE staff enabling this study by recording elephant life events and morphology for so long. Particularly, we thank Mu Mu Thein and Nina Aro for their patience in helping translating countless elephant morphological measurements from burmese to english. We also thank Carly Lynsdale for ensuring our english was fluent. We thank *the Academy of Finland, the European Research Council and the Ella & Georg Ehrnrooth Foundation for fundings*.

## Ethics

The study was performed with the permission from the Myanmar Ministry of Natural Resources and Environmental Conservation and following the University of Turku ethical guidelines.

## Conflict of interest

The authors have no conflict of interest to declare.

## Funding

This work was supported by the Academy of Finland [292368], the European Research Council [ERC-2014-CoG 648766] and the Ella & Georg Ehrnrooth Foundation.

## Authors’ contribution

HHA, WH and UKN performed field work and data collection. VL, VB and MB designed the study and LL selected, extracted and translated data. LL carried out all statistical analyses with contributions from VB and MB. LL wrote the manuscript and all authors critically reviewed it. All authors approved the manuscript for publication and agree to be held accountable for the content therein and approve the final version of the manuscript.

## Data accessibility

The datasets and codes supporting the conclusions of this article have been uploaded on Dryad digital repository and will be made publicly available upon acceptance.

## Notes

### Competing Interest Statement

The authors have declared no competing interest.

### Summary of Updates

Main changes concern: - Figures: The main manuscript include only the GLMM trajectories, while GAMM trajectories are presented in Supp (SI7, Fig. S4). X-axis is now in age (years) to facilitate interpretation for the reader, but the original figure with delta age for x-axis is available in supp (SI5, Fig. S2). - Discussion: We more extensively discuss the results found in the light of previous results of actuarial and reproductive ageing found in this population (particularly the idea of mosaic ageing in females).

## References

Andersen, R., Gaillard, J.-M., Linnell, J. D. C., & Duncan, P. (2000). Factors affecting maternal care in an income breeder, the European roe deer. Journal of Animal Ecology, 69(4), 672–682.

Bartoń, K. (2021). MuMln⍰: Multi-Model Inference (1.43.17) [Computer software]. https://CRAN.R-project.org/package=MuMIn

Bates, D., Mächler, M., Bolker, B., & Walker, S. (2015). Fitting linear mixed-effects models using lme4. Journal of Statistical Software, 67(1), 1–48. https://doi.org/10.18637/jss.v067.i01

Beirne, C., Delahay, R., & Young, A. (2015). Sex differences in senescence⍰: The role of intra-sexual competition in early adulthood. Proceedings of the Royal Society B: Biological Sciences, 282(1811), 20151086. https://doi.org/10.1098/rspb.2015.1086

Bérubé, C. H., Festa-Bianchet, M., & Jorgenson, J. T. (1999). Individual Differences, Longevity, and Reproductive Senescence in Bighorn Ewes. Ecology, 80(8), 2555–2565. https://doi.org/10.2307/177240

Bolker, B. M. (2008). Ecological Models and Data in R. Princeton University Press. https://press.princeton.edu/books/hardcover/9780691125220/ecological-models-and-data-in-r

Bonduriansky, R., Maklakov, A., Zajitschek, F., & Brooks, R. (2008). Sexual selection, sexual conflict and the evolution of ageing and life span. Functional Ecology, 22(3), 443–453. https://doi.org/10.1111/j.1365-2435.2008.01417.x

Briga, M., Jimeno, B., & Verhulst, S. (2019). Coupling lifespan and aging? The age at onset of body mass decline associates positively with sex-specific lifespan but negatively with environment-specific lifespan. Experimental Gerontology, 119, 111–119. https://doi.org/10.1016/j.exger.2019.01.030

Briga, M., & Verhulst, S. (2021). Mosaic metabolic ageing⍰: Basal and standard metabolic rate age in opposite directions and independent of environmental quality, sex and lifespan in a passerine. Functional Ecology, 35. https://doi.org/10.1111/1365-2435.13785

Bronikowski, A. M., Altmann, J., Brockman, D. K., Cords, M., Fedigan, L. M., Pusey, A., Stoinski, T., Morris, W. F., Strier, K. B., & Alberts, S. C. (2011). Aging in the Natural World⍰: Comparative Data Reveal Similar Mortality Patterns Across Primates. Science (New York, N.y.), 331(6022), 1325–1328. https://doi.org/10.1126/science.1201571

Brown, J. L. (2014). Comparative reproductive biology of elephants. Advances in Experimental Medicine and Biology, 753, 135–169. https://doi.org/10.1007/978-1-4939-0820-2_8

Burnham, K. P., Anderson, D. R., & Huyvaert, K. P. (2011). AIC model selection and multimodel inference in behavioral ecology⍰: Some background, observations, and comparisons. Behavioral Ecology and Sociobiology, 65(1), 23–35. https://doi.org/10.1007/s00265-010-1029-6

Camus, M. F., Clancy, D. J., & Dowling, D. K. (2012). Mitochondria, maternal inheritance, and male aging. Current Biology: CB, 22(18), 1717–1721. https://doi.org/10.1016/j.cub.2012.07.018

Carranza, J., & Pérez-Barbería, F. J. (2007). Sexual selection and senescence⍰: Male size-dimorphic ungulates evolved relatively smaller molars than females. The American Naturalist, 170(3), 370–380. https://doi.org/10.1086/519852

Chen, H., & Maklakov, A. A. (2014). Condition Dependence of Male Mortality Drives the Evolution of Sex Differences in Longevity. Current Biology, 24(20), 2423–2427. https://doi.org/10.1016/j.cub.2014.08.055

Clutton-Brock, T. H., & Isvaran, K. (2007). Sex differences in ageing in natural populations of vertebrates. Proceedings of the Royal Society B: Biological Sciences, 274(1629), 3097–3104. https://doi.org/10.1098/rspb.2007.1138

Coulson, J. C., & Fairweather, J. A. (2001). Reduced reproductive performance prior to death in the Black-legged Kittiwake⍰: Senescence or terminal illness? Journal of Avian Biology, 32(2), 146–152. https://doi.org/10.1034/j.1600-048X.2001.320207.x

Crawley, J. a. H., Mumby, H. S., Chapman, S. N., Lahdenperä, M., Mar, K. U., Htut, W., Soe, A. T., Aung, H. H., & Lummaa, V. (2017). Is bigger better? The relationship between size and reproduction in female Asian elephants. Journal of Evolutionary Biology, 30(10), 1836–1845. https://doi.org/10.1111/jeb.13143

Douhard, F., Gaillard, J.-M., Pellerin, M., Jacob, L., & Lemaître, J.-F. (2017). The cost of growing large⍰: Costs of post-weaning growth on body mass senescence in a wild mammal. Oikos, 126(9), 1329–1338. https://doi.org/10.1111/oik.04421

Eisenberg, J. F., McKay, G. M., & Jainudeen, M. R. (1971). Reproductive behavior of the Asiatic elephant (Elephas maximus maximus L.). Behaviour, 38(3), 193–225. https://doi.org/10.1163/156853971x00087

Flatt, T. (2005). The evolutionary genetics of canalization. The Quarterly Review of Biology, 80(3), 287–316. https://doi.org/10.1086/432265

Gaillard, J.-M., Berger, V., Tidière, M., Duncan, P., & Lemaître, J.-F. (2015). Does tooth wear influence ageing? A comparative study across large herbivores. Experimental Gerontology, 71, 48–55. https://doi.org/10.1016/j.exger.2015.09.008

Gaillard, J.-M., Festa-Bianchet, M., Delorme, D., & Jorgenson, J. (2000). Body mass and individual fitness in female ungulates⍰: Bigger is not always better. Proceedings of the Royal Society of London. Series B: Biological Sciences, 267(1442), 471–477. https://doi.org/10.1098/rspb.2000.1024

Global BMI Mortality Collaboration, Di Angelantonio, E., Bhupathiraju, S., Wormser, D., Gao, P., Kaptoge, S., Berrington de Gonzalez, A., Cairns, B., Huxley, R., Jackson, C., Joshy, G., Lewington, S., Manson, J., Murphy, N., Patel, A., Samet, J., Woodward, M., Zheng, W., Zhou, M.,… Hu, F. (2016). Body-mass index and all-cause mortality⍰: Individual-participant-data meta-analysis of 239 prospective studies in four continents. The Lancet, 388(10046), 776–786. https://doi.org/10.1016/S0140-6736(16)30175-1

Hämäläinen, A., Dammhahn, M., Aujard, F., Eberle, M., Hardy, I., Kappeler, P. M., Perret, M., Schliehe-Diecks, S., & Kraus, C. (2014). Senescence or selective disappearance? Age trajectories of body mass in wild and captive populations of a small-bodied primate. Proceedings of the Royal Society B: Biological Sciences, 281(1791), 20140830. https://doi.org/10.1098/rspb.2014.0830

Hayward, A. D., Mar, K. U., Lahdenperä, M., & Lummaa, V. (2014). Early reproductive investment, senescence and lifetime reproductive success in female Asian elephants. Journal of Evolutionary Biology, 27(4), 772–783. https://doi.org/10.1111/jeb.12350

Hayward, A. D., Moorad, J., Regan, C. E., Berenos, C., Pilkington, J. G., Pemberton, J. M., & Nussey, D. H. (2015). Asynchrony of senescence among phenotypic traits in a wild mammal population. Experimental Gerontology, 71, 56–68. https://doi.org/10.1016/j.exger.2015.08.003

Jones, O. R., Scheuerlein, A., Salguero-Gómez, R., Camarda, C. G., Schaible, R., Casper, B. B., Dahlgren, J. P., Ehrlén, J., García, M. B., Menges, E. S., Quintana-Ascencio, P. F., Caswell, H., Baudisch, A., & Vaupel, J. W. (2014). Diversity of ageing across the tree of life. Nature, 505(7482), 169–173. https://doi.org/10.1038/nature12789

Klemera, P., & Doubal, S. (2006). A new approach to the concept and computation of biological age. Mechanisms of Ageing and Development, 127(3), 240–248. https://doi.org/10.1016/j.mad.2005.10.004

Lahdenperä, M., Mar, K. U., Courtiol, A., & Lummaa, V. (2018). Differences in age-specific mortality between wild-caught and captive-born Asian elephants. Nature Communications, 9(1), 3023. https://doi.org/10.1038/s41467-018-05515-8

Lahdenperä, M., Mar, K. U., & Lummaa, V. (2016). Nearby grandmother enhances calf survival and reproduction in Asian elephants. Scientific Reports, 6(1), 27213. https://doi.org/10.1038/srep27213

Leimgruber, P., Senior, B., Aung, M., Songer, M., Mueller, T., Wemmer, C., & Ballou, J. (2008). Modeling population viability of captive elephants in Myanmar (Burma)⍰: Implications for wild populations. Animal Conservation, 11, 198–205. https://doi.org/10.1111/j.1469-1795.2008.00172.x

Lemaître, J.-F., Ronget, V., Tidière, M., Allainé, D., Berger, V., Cohas, A., Colchero, F., Conde, D. A., Garratt, M., Liker, A., Marais, G. A. B., Scheuerlein, A., Székely, T., & Gaillard, J.-M. (2020). Sex differences in adult lifespan and aging rates of mortality across wild mammals. Proceedings of the National Academy of Sciences, 117(15), 8546–8553. https://doi.org/10.1073/pnas.1911999117

Lynch, E. C., Lummaa, V., Htut, W., & Lahdenperä, M. (2019). Evolutionary significance of maternal kinship in a long-lived mammal. Philosophical Transactions of the Royal Society B: Biological Sciences, 374(1780), 20180067. https://doi.org/10.1098/rstb.2018.0067

Maklakov, A. A., & Lummaa, V. (2013). Evolution of sex differences in lifespan and aging⍰: Causes and constraints. BioEssays, 35, 717–724. https://doi.org/10.1002/bies.201300021

Monaghan, P., Charmantier, A., Nussey, D. H., & Ricklefs, R. E. (2008). The evolutionary ecology of senescence. Functional Ecology, 22(3), 371–378. https://doi.org/10.1111/j.1365-2435.2008.01418.x

Mumby, H. S., Chapman, S. N., Crawley, J. A. H., Mar, K. U., Htut, W., Thura Soe, A., Aung, H. H., & Lummaa, V. (2015). Distinguishing between determinate and indeterminate growth in a long-lived mammal. BMC Evolutionary Biology, 15(1), 214. https://doi.org/10.1186/s12862-015-0487-x

Mumby, H. S., Mar, K. U., Hayward, A. D., Htut, W., Htut-Aung, Y., & Lummaa, V. (2015). Elephants born in the high stress season have faster reproductive ageing. Scientific Reports, 5(1), 13946. https://doi.org/10.1038/srep13946

Mumby, H. S., Mar, K. U., Thitaram, C., Courtiol, A., Towiboon, P., Min-Oo, Z., Htut-Aung, Y., Brown, J. L., & Lummaa, V. (2015). Stress and body condition are associated with climate and demography in Asian elephants. Conservation Physiology, 3(1), cov030. https://doi.org/10.1093/conphys/cov030

Nussey, D. H., Coulson, T., Delorme, D., Clutton-Brock, T. H., Pemberton, J. M., Festa-Bianchet, M., & Gaillard, J.-M. (2011). Patterns of body mass senescence and selective disappearance differ among three species of free-living ungulates. Ecology, 92(10), 1936–1947. JSTOR. https://doi.org/10.1890/11-0308.1

Nussey, D. H., Kruuk, L., Morris, A., Clements, M., Pemberton, J., & Clutton-Brock, T. (2009). Inter- and Intrasexual Variation in Aging Patterns across Reproductive Traits in a Wild Red Deer Population. The American Naturalist, 174, 342–357. https://doi.org/10.1086/603615

Oo, Z. M. (2010). The training methods used in Myanma Timber Enterprise. Gajah, 33, 58–61.

Pelletier, F., Clutton-Brock, T., Pemberton, J., Tuljapurkar, S., & Coulson, T. (2007). The Evolutionary Demography of Ecological Change⍰: Linking Trait Variation and Population Growth. Science, 315(5818), 1571–1574. https://doi.org/10.1126/science.1139024

Prospective Studies Collaboration. (2009). Body-mass index and cause-specific mortality in 900 000 adults⍰: Collaborative analyses of 57 prospective studies. The Lancet, 373(9669), 1083–1096. https://doi.org/10.1016/S0140-6736(09)60318-4

R Core Team. (2021). R: A language and environment for statistical computing. R Foundation for Statistical Computing. https://www.R-project.org

Rando, T. A., & Wyss-Coray, T. (2021). Asynchronous, contagious and digital aging. Nature Aging, 1(1), 29–35. https://doi.org/10.1038/s43587-020-00015-1

Reichert, S., Berger, V., dos Santos, D. J. F., Lahdenperä, M., Nyein, U. K., Htut, W., & Lummaa, V. (2022). Age related variation of health markers in Asian elephants. Experimental Gerontology, 157, 111629. https://doi.org/10.1016/j.exger.2021.111629

Robinson, M. R., Mar, K. U., & Lummaa, V. (2012). Senescence and age-specific trade-offs between reproduction and survival in female Asian elephants⍰: Age-specific reproduction and survival. Ecology Letters, 15(3), 260–266. https://doi.org/10.1111/j.1461-0248.2011.01735.x

Seltmann, M. W., Helle, S., Htut, W., & Lahdenperä, M. (2019). Males have more aggressive and less sociable personalities than females in semi-captive Asian elephants. Scientific Reports, 9(1), 2668. https://doi.org/10.1038/s41598-019-39915-7

Simons, M. J. P., Briga, M., & Verhulst, S. (2016). Stabilizing survival selection on presenescent expression of a sexual ornament followed by a terminal decline. Journal of Evolutionary Biology, 29(7), 1368–1378. https://doi.org/10.1111/jeb.12877

Simpson, G. L., & Singmann, H. (2021). gratia⍰: Graceful ‘ggplot’-based graphics and other functions for GAMs fitted using « mgcv » (0.6.0) [Computer software].

Sukumar, R. (2003). The living elephants⍰: Evolutionary ecology, behavior, and conservation. Oxford University Press.

Tafani, M., Cohas, A., Bonenfant, C., Gaillard, J.-M., Lardy, S., & Allainé, D. (2013). Sex-specific senescence in body mass of a monogamous and monomorphic mammal⍰: The case of Alpine marmots. Oecologia, 172(2), 427–436. https://doi.org/10.1007/s00442-012-2499-1

Thorley, J., Duncan, C., Sharp, S. P., Gaynor, D., Manser, M. B., & Clutton-Brock, T. (2020). Sex-independent senescence in a cooperatively breeding mammal. Journal of Animal Ecology, 89(4), 1080–1093. https://doi.org/10.1111/1365-2656.13173

Tidière, M., Gaillard, J.-M., Müller, D. W. H., Lackey, L. B., Gimenez, O., Clauss, M., & Lemaître, J.-F. (2015). Does sexual selection shape sex differences in longevity and senescence patterns across vertebrates? A review and new insights from captive ruminants. Evolution, 69(12), 3123–3140. https://doi.org/10.1111/evo.12801

Toke Gale, U. (1974). Burmese timber elephant. Trade Corporation.

Tower, J. (2006). Sex-specific regulation of aging and apoptosis. Mechanisms of Ageing and Development, 127(9), 750–718. https://doi.org/10.1016/j.mad.2006.05.001

van de Pol, M., & Verhulst, S. (2006). Age-dependent traits⍰: A new statistical model to separate within- and between-individual effects. The American Naturalist, 167(5), 766–773. https://doi.org/10.1086/503331

van de Pol, M., & Wright, J. (2009). A simple method for distinguishing within-versus between-subject effects using mixed models. Animal Behaviour, 77(3), 753–758. https://doi.org/10.1016/j.anbehav.2008.11.006

Walker, L. C., & Herndon, J. G. (2010). Mosaic aging. Medical Hypotheses, 74(6), 1048–1051. https://doi.org/10.1016/j.mehy.2009.12.031

Williams, G. C. (1957). Pleiotropy, natural selection, and the evolution of senescence. Evolution, 11(4), 398–411. https://doi.org/10.2307/2406060

Wood, S. N. (2011). Fast stable restricted maximum likelihood and marginal likelihood estimation of semiparametric generalized linear models. Journal of the Royal Statistical Society (B), 73(1), 3–36. https://doi.org/10.1111/j.1467-9868.2010.00749.x

Zaw, U. K. (1997). Utilization of elephants in timber harvesting in Myanmar. Gajah, 17, 9–22.

Zuur, A., leno, E. N., Walker, N., Saveliev, A. A., & Smith, G. M. (2009). Mixed Effects Models and Extensions in Ecology with R. Springer-Verlag. https://doi.org/10.1007/978-0-387-87458-6

