## Supplementary Information for "Sex-specific body mass ageing trajectories in adult Asian elephants"

### Authors informations

Lucas D. Lalande[1,2,3,a] (ORCID iD: 0000-0001-9319-8033),

Virpi Lummaa[1,a] (ORCID iD: 0000-0002-2128-7587),

Htoo H. Aung[4],

Win Htut[4],

U. Kyaw Nyein[4],

Vérane Berger[1,a,*] (ORCID iD: 0000-0002-2153-0719),

Michael Briga[1,a,*] (ORCID iD: 0000-0003-3160-0407)

[1] Department of Biology, University of Turku, Turku, Finland;

[2] Université Bourgogne Franche-Comté, Dijon, France;

[3] Present address: Université de Lyon, Université Lyon 1, UMR CNRS 5558, Villeurbanne CEDEX, France;

[4] Myanma Timber Enterprise, Ministry of Natural Resources and Environmental Conservation, West Gyogone Forest Compound, Yangon, Myanmar;

[*] Equal contribution of these authors

### Supplementary Information 1. Body mass estimation

Because of the high correlation of chest girth (CG) with body mass, compared to other morphological metrics (Chapman et al., 2016), we performed a model selection according to AICc values (Akaike Information Criteria corrected) to choose the best way of estimating body masses from CG for males and females separately, as they have different weight growth curves (Chapman et al., 2016; Mumby, Chapman, et al., 2015). In addition to CG being a good predictor of body mass, we also included height to shoulders (H) in the models as indicative of structural size in Asian elephants. Our data indicate a robust correlation between CG and body mass for individuals with both measurements ($r_{males}$=0.80; $r_{females}$=0.71). The correlation between CG and body mass is higher than the correlation between CG and H (*i.e.* structural size ($r_{males}$=0.64; $r_{females}$=0.51). Body mass estimations and equations were estimated comparing n = 1,470 ($n_{male}$ = 491, $n_{female}$ = 979) known body mass measurements to the same number of both CG and H measurements.

For females, the full model, including both CG and H and their quadratic effects, showed lower AICc and the best correlation between estimations and measurements, while it was the model including the linear and quadratic effect of CG, and the linear effect of H which was retained for males (table S1). We would rather use the most accurate equations to predict body mass from other metrics, although CG and H were correlated for both sexes ($r_{males}$ = 0.65, t = 18.7, p < 0.0001; $r_{females}$ = 0.51, t = 18.5, p < 0.0001). From the coefficients of the selected model, we formulated equations (1) and (2) for males and females respectively. The last term of both equations was added to correct for the tendency of the equations to overestimate body masses. Correlations between estimated body masses from equations (1) and (2) and known body masses was r = 0.90 [95%CI: 0.89; 0.91].

#### Males

$$BM = 2{,}829 - 32.17 \times CG + 0.06 \times CG^2 + 17.57 \times H - 63.26 \quad (1)$$

#### Females

$$BM = 7{,}697 - 16.40 \times CG + 0.04 \times CG^2 - 48.77 \times H + 0.14 \times H^2 - 21.26 \quad (2)$$

**Table S1.** Best predictors (in bold) of body mass estimation based on morphometric measures based on the second-order Akaike information criterion (AICc). CG: chest girth, H: height, r: spearman correlation coefficient, k: degrees of freedom.

| Response | Model | r | Male | | | Female | | |
|---|---|---|---|---|---|---|---|---|
| | | | AICc | k | r | AICc | k | r |
| Body mass | CG | 0.851 | 7,116.7 | 3 | 0.800 | 13,596.5 | 3 | 0.711 |
| | CG + $CG^2$ | 0.858 | 7,096.8 | 4 | 0.810 | 13,559.0 | 4 | 0.725 |
| | CG + H | 0.895 | 6,849.4 | 4 | 0.890 | 13,434.9 | 4 | 0.763 |
| | **CG + $CG^2$ + H** | **0.902** | **6,802.2** | **5** | **0.901** | 13,404.2 | 5 | 0.772 |
| | CG + H + $H^2$ | 0.898 | 6,841.5 | 5 | 0.892 | 13,406.0 | 5 | 0.771 |
| | **CG + $CG^2$ + H + $H^2$** | -0.461 | 6,803.4 | 6 | 0.496 | **13,389.8** | **6** | **0.776** |

### Supplementary Information 2. Illustration of body mass ageing trajectories using GLMMs

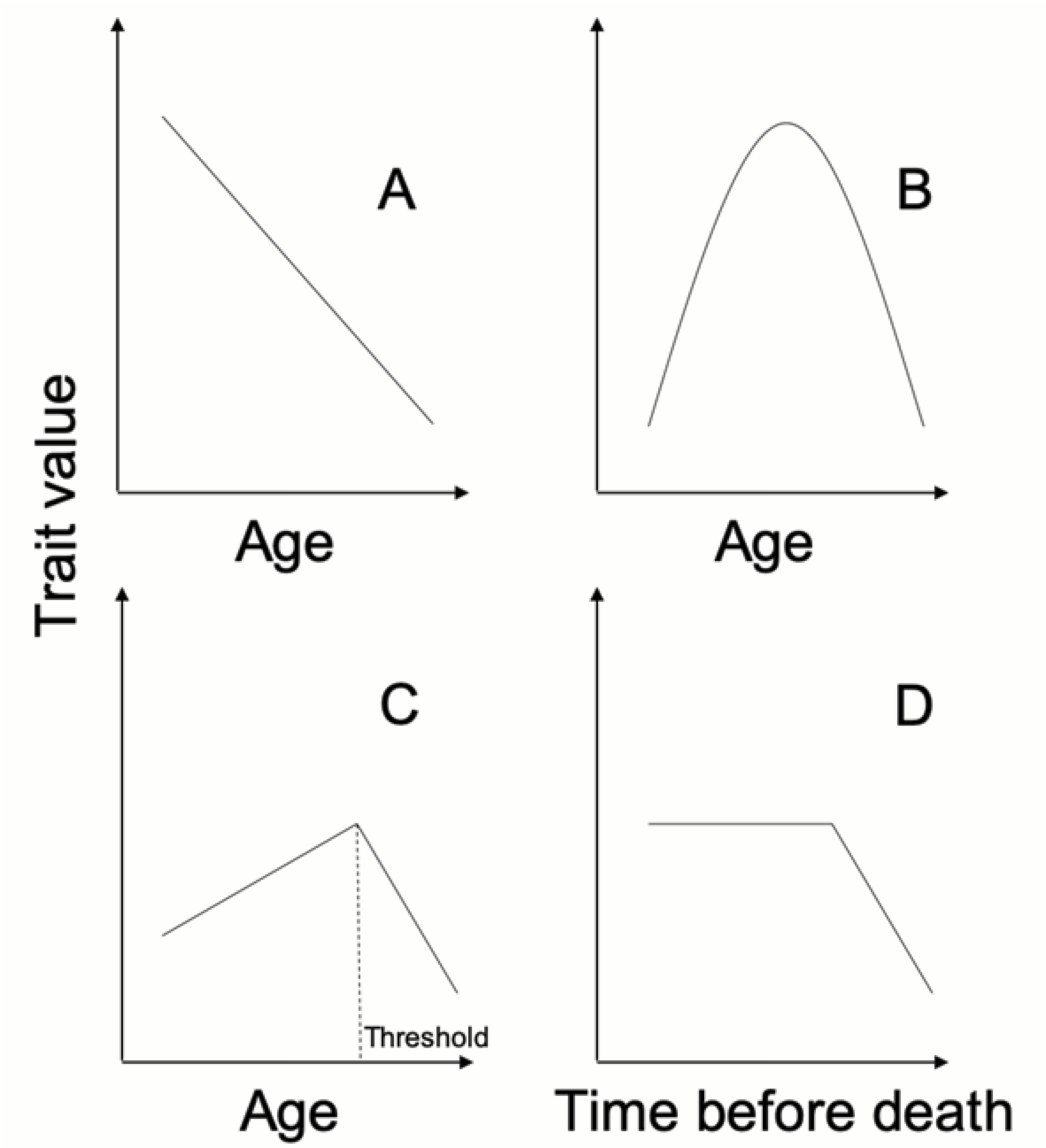

**Figure S1.** Schematic representation of the four ageing trajectories tested. Using the individual's chronological age, ageing trajectory can be (A) linear, (B) quadratic, or (C) with a threshold. However, using the time before death, ageing can correspond to (D) a terminal decline. The slopes here are illustrative only and can, in practice, take any value. Figure adapted from Briga et al., 2019.

### Supplementary Information 3. Sex-specific age trajectories

**Table S2.** Evidence for sex-specific body mass ageing trajectories both in GAM and GLM models based on model selection approaches. The models are ranked from the best-fitting (lowest AICc, on the top) to poorer-fitting ones (higher AICc, at the bottom). Best-fitting in bold. AICc: second-order Akaike Information Criteria, ΔAICc: difference in AICc relative to the best model, k: degrees of freedom, w: model weight, s=smooth. All models contain the random intercepts individual identity *(1|ID)* and township *(1|township).*

| Model | Model | k | AICc | ΔAICc | w |
|---|---|---|---|---|---|
| GAMM | **log(bm) ~ terminal + sex + s(Δage) + age-last + age-last² + s(Δage, by=sex)** | **17.9** | **-8,809.1** | **0.0** | **1.0** |
| | log(bm) ~ terminal + sex + s(Δage) + Δage² + age-last + age-last² | 10.3 | -8,743.4 | 65.7 | 0.0 |
| GLMM | **log(bm) ~ terminal + sex + Δage + age-last + age-last² + sex:Δage** | **10.0** | **-8,730.6** | **0.0** | **1.0** |
| | log(bm) ~ terminal + sex + Δage + age-last + age-last² | 9.0 | -8,683.6 | 47.0 | 0.0 |

### Supplementary Information 4. Seasonal and spatial confounding variables

**Table S3.** Model selection of covariates based on the best fitting models in Table 1 describing body mass ageing trajectories for both sexes. The column 'Model' refers to the model names as given in Table 1. AICc: Akaike Information Criteria (corrected) of the selected models, k: degrees of freedom, 'season': ΔAICc (*i.e.* AICc differential compared to the selected model) when including the season of the body mass measurement, 'alive': ΔAICc when including whether individuals were dead or alive at the moment of the analysis, 'cw': ΔAICc when including whether the individuals were captive-born or wild-caught, 'measure': ΔAICc when including whether the body masses were estimated or measured.

| Sex | Model selected | AICc | ΔAICc | k | w |
|---|---|---|---|---|---|
| Males GAMMs | log(bm) ~ s(Δage) + age-last + cw | -3,222.8 | 0.0 | 9.1 | 0.234 |
| | **log(bm) ~ s(Δage) + age-last** | **-3,221.9** | **0.9** | **8.1** | **0.148** |
| | log(bm) ~ s(Δage) + age-last + cw + season | -3,221.2 | 1.6 | 11.1 | 0.104 |
| | log(bm) ~ s(Δage) + age-last + cw + measure | -3,220.8 | 2.0 | 10.1 | 0.085 |
| | log(bm) ~ s(Δage) + age-last + cw + alive | -3,220.4 | 2.4 | 11.1 | 0.071 |
| | log(bm) ~ s(Δage) + age-last + season | -3,220.2 | 2.6 | 10.1 | 0.066 |
| | log(bm) ~ s(Δage) + age-last + alive | -3,219.9 | 2.9 | 10.1 | 0.055 |
| | log(bm) ~ s(Δage) + age-last + measure | -3,219.8 | 2.9 | 9.1 | 0.054 |
| | log(bm) ~ s(Δage) + age-last + cw + measure + season | -3,219.1 | 3.6 | 12.1 | 0.038 |
| | log(bm) ~ s(Δage) + age-last + alive + measure + season | -3,218.7 | 4.1 | 13.1 | 0.031 |
| Females GAMMs | log(bm) ~ terminal + s(Δage) + age-last + age-last$^2$ + season | -5,641.5 | 0.0 | 12.8 | 0.316 |
| | log(bm) ~ terminal + s(Δage) + age-last + age-last$^2$ + season + measure | -5,640.2 | 1.4 | 13.7 | 0.160 |
| | **log(bm) ~ terminal + s(Δage) + age-last + age-last$^2$** | **-5,639.7** | **1.8** | **10.9** | **0.127** |
| | log(bm) ~ terminal + s(Δage) + age-last + age-last$^2$ + season + cw | -5,638.8 | 2.7 | 14.8 | 0.082 |
| | log(bm) ~ terminal + s(Δage) + age-last + age-last$^2$ + measure | -5,638.4 | 3.1 | 11.7 | 0.066 |
| | log(bm) ~ terminal + s(Δage) + age-last + age-last$^2$ + season + alive | -5,638.1 | 3.4 | 14.8 | 0.057 |
| | log(bm) ~ terminal + s(Δage) + age-last + age-last$^2$ + season + measure + cw | -5,637.5 | 4.1 | 15.9 | 0.042 |
| | log(bm) ~ terminal + s(Δage) + age-last + age-last$^2$ + cw | -5,637.0 | 4.5 | 12.9 | 0.033 |
| Males GLMMs | **log(bm) ~ Δage1 + Δage2 + age-last** | **-3,192.5** | **0.0** | **8.0** | **0.889** |
| | log(bm) ~ Δage1 + Δage2 + age-last + cw | -3,187.9 | 4.6 | 9.0 | 0.089 |
| | log(bm) ~ Δage1 + Δage2 + age-last + alive | -3,184.4 | 8.1 | 10.0 | 0.016 |
| Females GLMMs | **log(bm) ~ terminal + Δage + age-last + age-last$^2$** | **-5,603.6** | **0.0** | **8.0** | **0.983** |
| | log(bm) ~ terminal + Δage + age-last + age-last$^2$ + measure | -5,594.6 | 9.0 | 9.0 | 0.011 |

### Supplementary Information 5. Body mass ageing trajectories using GLMMs

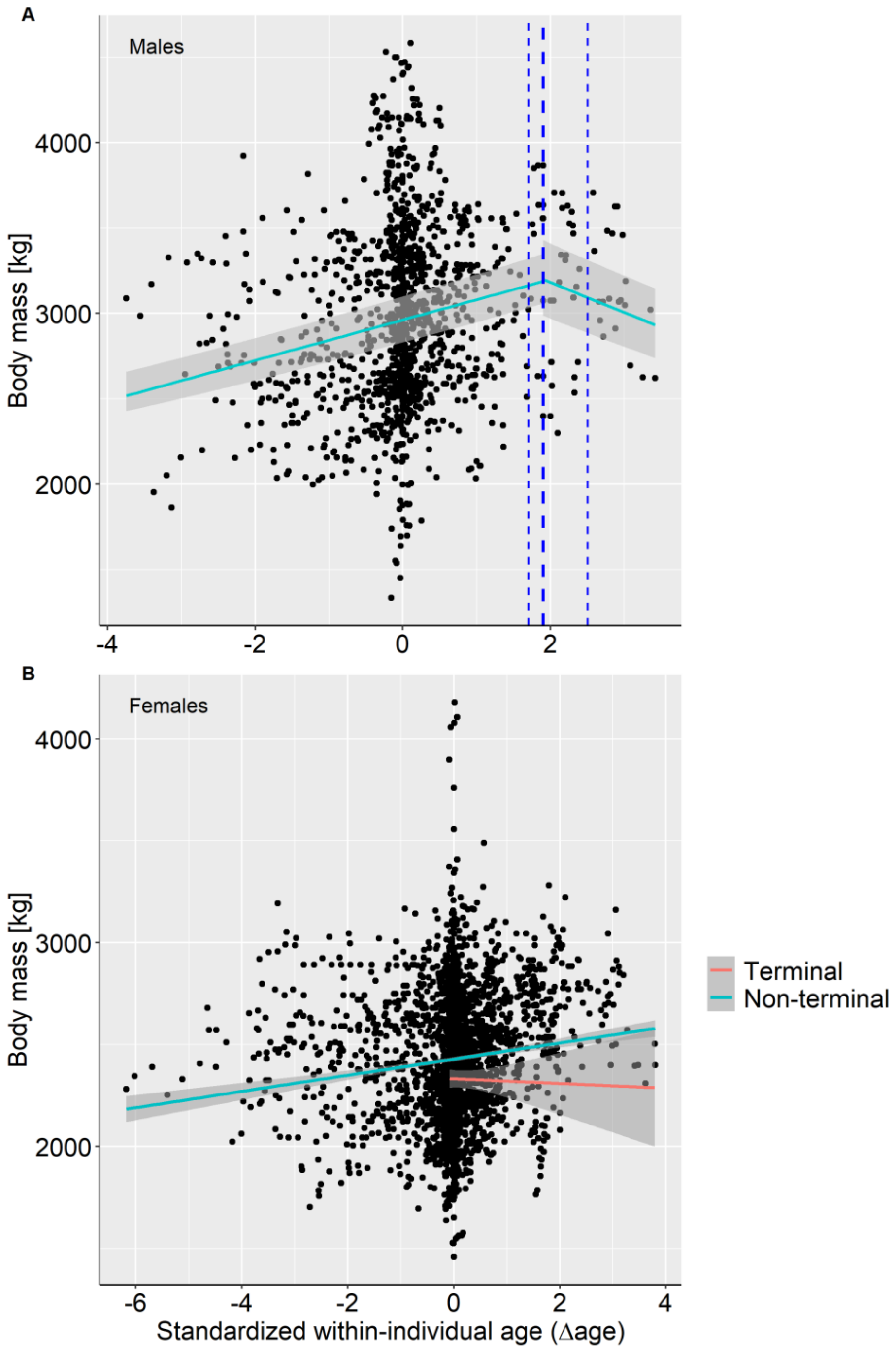

**Figure S2.** Body mass ageing trajectories of (A) males (n = 1,316 measurements on 171 individuals) and (B) females (n = 2,570 measurements on 322 individuals) with predictions of the best-fitting GLMMs (Table 1) with grey areas 95%CI. For males, the thick dashed-line shows the threshold age at onset of the body mass decline (1.9 or 48.3 years) with thin dashed-lines the 4 ΔAICc-CI [46.6, 52.3]. For females, measurements in the terminal year (red) are significantly lower (intercept) than measurements at other ages (blue). Note that the terminal slope is for illustration purposes only and was not statistically tested.

### Supplementary Information 6. Testing terminal decline windows

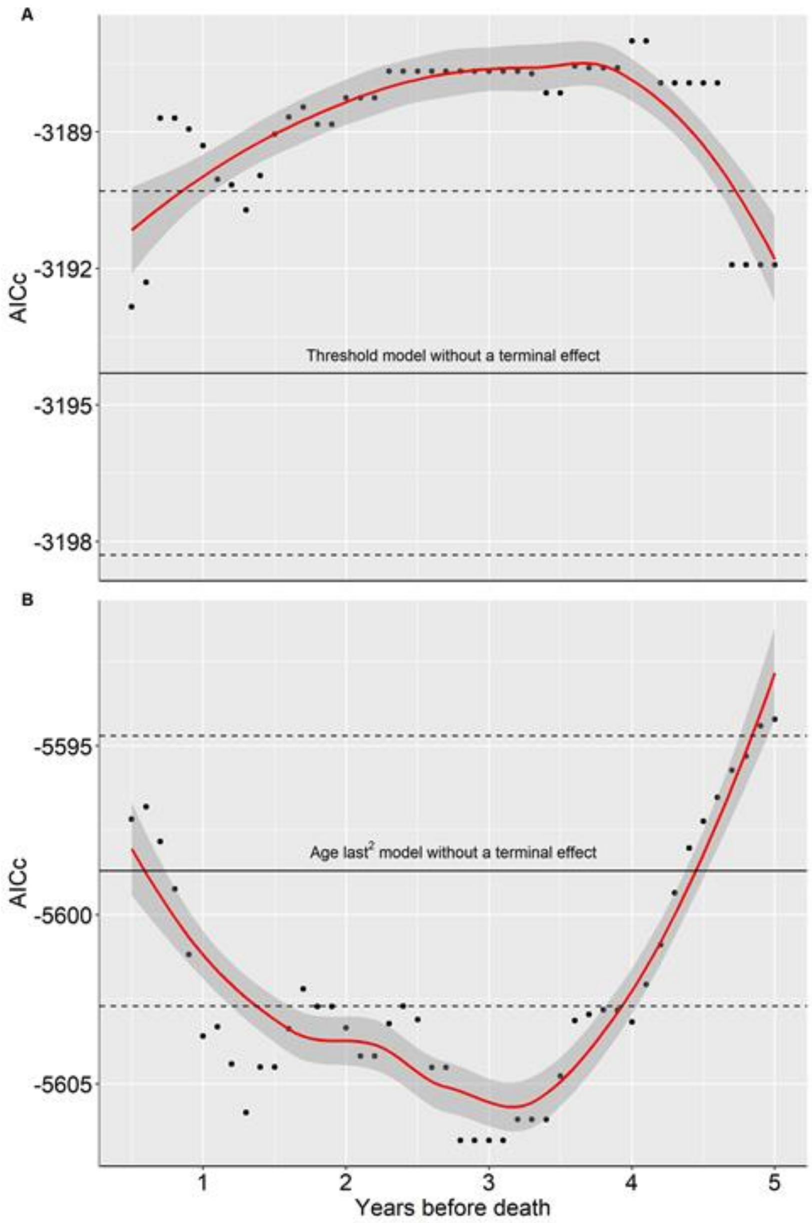

**Figure S3.** Testing variation in windows for the age at onset of the terminal decline for A) males and B) females. The black solid line represents the fit of the model selected without a terminal effect, and the dashed lines represent the confidence interval of this model (±4AICc). A) In males, terminal declines were not statistically supported (best fitting model in table 1), irrespective of the window used. B) In females, a statistically significant terminal decline occurs between 1 and 4 years before death and the statistical support through this windows is equivalent. Hence, we chose to illustrate models with terminal windows of one year.

### Supplementary Information 7. Body mass ageing trajectories using GAMMs

**Table S4.** Model selection of body mass ageing trajectories (bold) for males and females, using GAMMs for each model ageing trajectories ranked from the least to the most complex. AICc: second-order Akaike Information Criterion; ΔAICc: change in AICc relative to the best fitting model; k: degrees of freedom.

| Model type | Model | Males | | | Females | | |
|---|---|---|---|---|---|---|---|
| | | k | AICc | ΔAICc | k | AICc | ΔAICc |
| null | log(bm) ~ 1 | 4.0 | -2,835.8 | 386.1 | 4.0 | -5,422.6 | 217.1 |
| **smooth1** | **log(bm) ~ s(Δage) + age-last** | **8.1** | **-3,221.9** | **0.01** | 8.7 | -5,613.9 | 25.8 |
| +terminal | log(bm) ~ s(Δage) + age-last + terminal | 9.1 | -3,221.9 | 0.0 | 10.0 | -5,627.3 | 12.4 |
| smooth2 | log(bm) ~ s(Δage) + age-last + age-last$^2$ | 9.1 | -3,221.2 | 0.7 | 9.7 | -5,627.5 | 12.2 |
| **+terminal** | **log(bm) ~ s(Δage) + age-last + age-last$^2$ + terminal** | 10.1 | -3,221.3 | 0.6 | **10.9** | **-5,639.7** | **0.0** |

**Table S5.** Estimates of general additive mixed models (GAMMs) including individual body mass beyond 18 years of age as the response variable (in kg, log-transformed) for male and female Asian elephants. V: variance, SD: standard deviation, SE: standard-error, Df: degrees of freedom, F: Fisher value. Marginal and conditional $R^2$ give the variance explained by fixed effects, and both fixed and random effects, respectively.

| Males | | | Females | | |
|---|---|---|---|---|---|
| **Random effects** | **V** | **SD** | **Random effects** | **V** | **SD** |
| Individual identity | 0.018 | 0.136 | Individual identity | 0.009 | 0.095 |
| Township | 0.0004 | 0.021 | Township | 0.004 | 0.060 |
| **Fixed effects** | **Estimate** | **SE** | **Fixed effects** | **Estimate** | **SE** |
| Intercept | 7.997 | 0.013 | Intercept | 7.832 | 0.009 |
| Age at last measurement | 0.089 | 0.012 | Age at last measurement | 0.034 | 0.007 |
| | | | Age at last measurement$^2$ | -0.023 | 0.006 |
| | | | Terminal (1) | -0.075 | 0.020 |
| Smooth term | Df | F | Smooth term | Df | F |
| Δage | 6.096 | 70.4 | Δage | 6.899 | 27.9 |
| **Marginal $R^2$** | 0.23 | | **Marginal $R^2$** | 0.10 | |
| **Conditional $R^2$** | 0.89 | | **Conditional $R^2$** | 0.80 | |

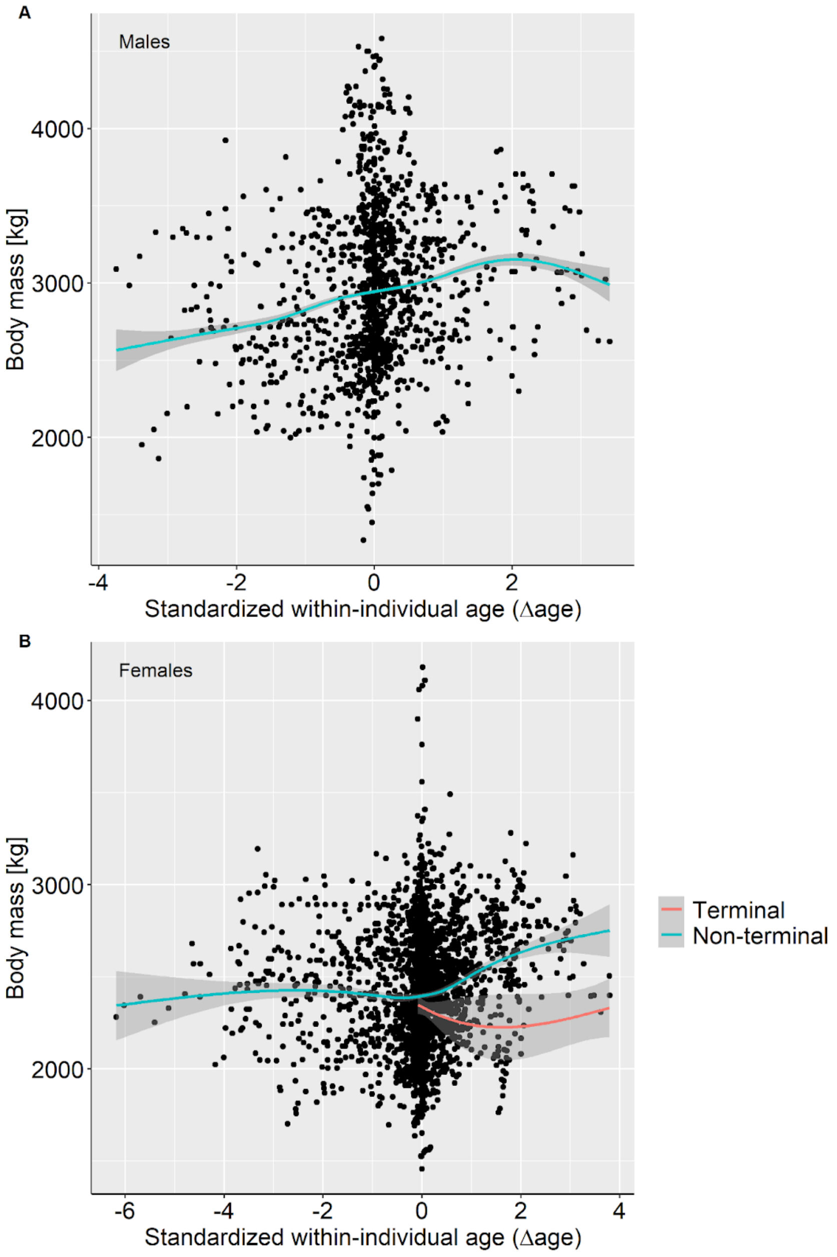

**Figure S4.** Body mass ageing trajectories of (A) males (n=1,316 measurements on 171 individuals) and (B) females (n=2,570 measurements on 322 individuals) with solid lines showing predictions of the best-fitting GAM models (Table S4) and grey areas 95%CI. For females, measurements in the terminal year (red) are significantly lower than measurements at other ages (grey), but note that the association (slope) with Δage is for illustration purposes only and was not statistically tested.

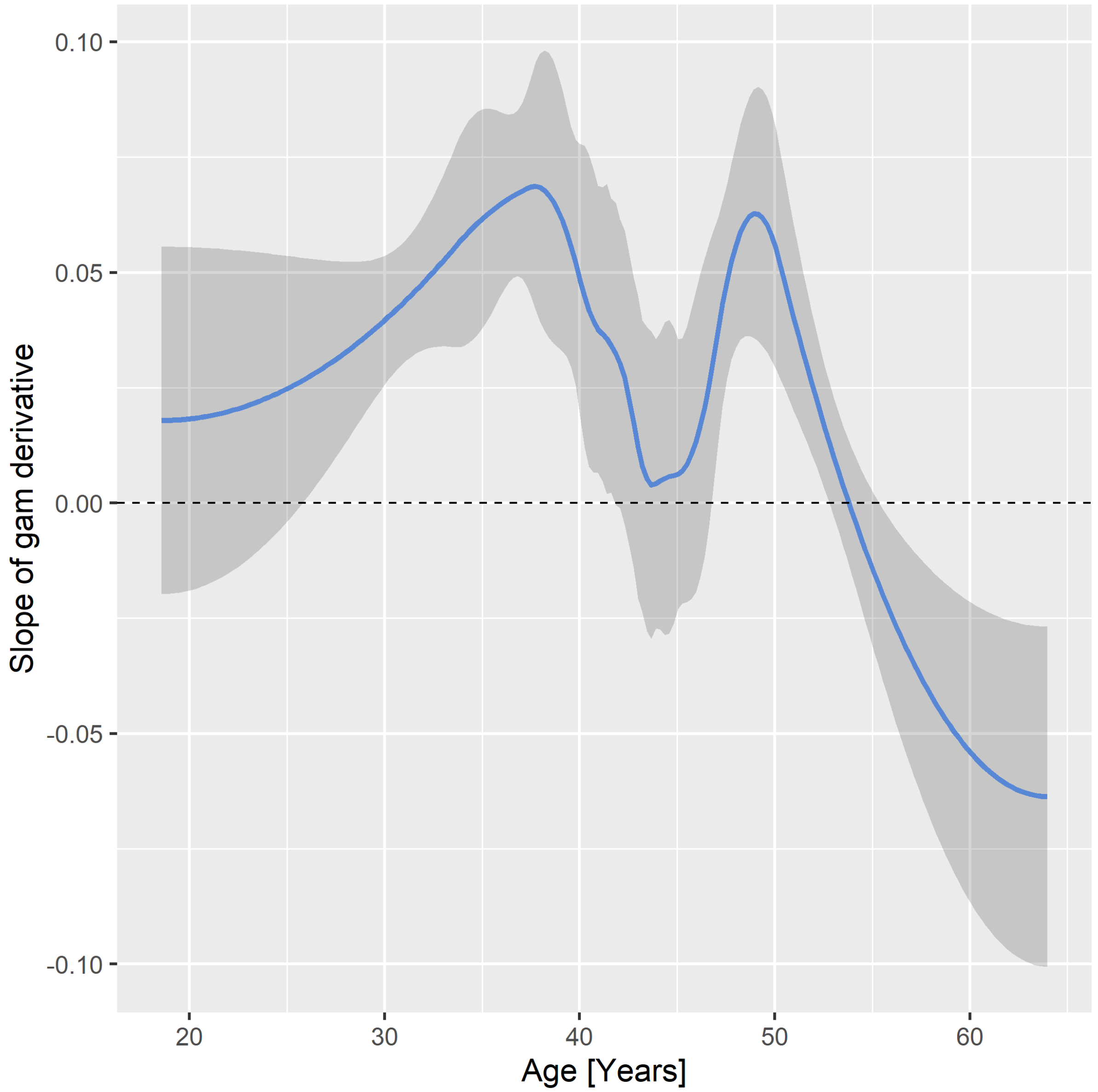

**Figure S5.** Derivative of the best fitting GAM model in males (Table S4, smooth 1) to identify the age at which maximum body mass is reached, *i.e.* when the derivative (blue line) is zero. Grey zones indicate the 95% confidence interval around the age at maximum.

### References

Briga, M., Jimeno, B., & Verhulst, S. (2019). Coupling lifespan and aging? The age at onset of body mass decline associates positively with sex-specific lifespan but negatively with environment-specific lifespan. *Experimental Gerontology*, *119*, 111-119. https://doi.org/10.1016/j.exger.2019.01.030

Chapman, S. N., Mumby, H. S., Crawley, J. A. H., Mar, K. U., Htut, W., Soe, A. T., Aung, H. H., & Lummaa, V. (2016). How big is it really? Assessing the efficacy of indirect estimates of body size in Asian elephants. *PLOS ONE*, *11*(3), e0150533. https://doi.org/10.1371/journal.pone.0150533

Mumby, H. S., Chapman, S. N., Crawley, J. A. H., Mar, K. U., Htut, W., Thura Soe, A., Aung, H. H., & Lummaa, V. (2015). Distinguishing between determinate and indeterminate growth in a long-lived mammal. *BMC Evolutionary Biology*, *15*(1), 214. https://doi.org/10.1186/s12862-015-0487-x
